# Neural responses to reconstructed target pursuits

**DOI:** 10.1101/2023.05.03.539331

**Authors:** Yuri Ogawa, Sarah Nicholas, Malin Thyselius, Richard Leibbrandt, Thomas Nowotny, James C. Knight, Karin Nordström

## Abstract

Many animals use motion vision information to control dynamic behaviors. Predatory animals, for example, show an exquisite ability to detect rapidly moving prey followed by pursuit and capture. Such target detection is not only used by predators but can also play an important role in conspecific interactions. Male hoverflies (*Eristalis tenax*), for example, vigorously defend their territories against conspecific intruders. Visual target detection is believed to be subserved by specialized target tuned neurons that are found in a range of species, including cats, zebrafish, and insects. However, how these target-tuned neurons respond to actual pursuit trajectories is currently not well understood. To redress his, we recorded extracellularly from target selective descending neurons (TSDNs) in male *Eristalis tenax* hoverflies. We show that the neurons have dorso-frontal receptive fields, with a preferred direction up and away from the visual midline. We next reconstructed visual flow-fields as experienced during pursuits of artificial targets (black beads). We recorded TSDN responses to six reconstructed pursuits and found that each neuron responded consistently at remarkably specific time points, but that these time points differed between neurons. We compared the observed spike probability with the spike probability predicted from each neuron’s receptive field and size tuning, and found a correlation coefficient of 0.35. Interestingly, however, the overall response rate was low, with individual neurons responding to only a small part of each reconstructed pursuit. In contrast, the TSDN population responded to a substantially larger proportion of the pursuits (up to a median of 23%). This large variation between neurons could be useful if different neurons control different parts of the behavioral output.

**Significance statement:** Descending neurons constitute less than 1% of the nervous system, yet have to convey all requisite information from the brain to the body. They are therefore a crucial bottleneck in sensorimotor transformation. Descending target tuned neurons in male hoverflies (*Eristalis tenax)*, for example, are believed to play a key role in territory defense and pursuit of conspecifics. However, this has not been tested using visual stimuli resembling reconstructed target pursuits. We here found that the observed neural responses to reconstructed pursuits are stronger than those predicted from responses to simpler stimuli. In addition, while the responses to simple stimuli suggested a homogenous population of neurons, the reconstructed pursuits showed important differences between individual neurons. Our data thus highlight the need for using more naturalistic stimuli when deciphering neural function.

## Introduction

Many animals use motion information to control fast and dynamic behaviors. For example, predators rely on sensory input when identifying and pursuing small, fast-moving prey, such as bats using echolocation^1^ and zebrafish larvae using visual information^2, 3^. Many insects, including dragonflies and robber flies, are also excellent predators that use visual information to locate their small, moving prey^4, 5^. Importantly, however, similar target detection behaviors may also be used in conspecific interactions. Male hoverflies (*Eristalis tenax*), for example, use visual cues to vigorously guard their territories, followed by high-speed pursuit of any territorial intruders, or courtship interactions with potential mates^6^.

The neural circuits that support such target detection behaviors have been described in a range of species, including the salamander retina^7^, the bat inferior colliculus^8^, the tectum and nucleus isthmi of zebrafish larvae^9,10^, and the optic lobes of dragonflies^11^ and hoverflies^12^. Indeed, these sensory pathways appear to be optimally tuned to the unique parameters of moving prey.

Visual neurons that are hypothesized to be involved in target pursuit exhibit size and speed tuning to abstract visual stimuli that are similar to the images likely experienced during pursuit (for review, see e.g. Ref. ^13^). However, these experiments often used target stimuli with constant velocities and linear paths, and it is thus unclear how the visual neurons respond to more naturalistic target trajectories that change dynamically^14, 15^. This is important as many sensory systems are known to respond differently to naturalistic input compared with more restricted stimuli. For example, in response to naturalistic input neurons in the bat inferior colliculus show higher signal-to-noise ratio and improved parallel processing^8^. In rodents, individual primary sensory neurons responding to tactile whisker information simultaneously encode multi-dimensional components when exposed to naturalistic stimuli, not expected from their responses to more restricted stimuli^16^.

In insects, target pursuit is likely subserved by specialized target-tuned neurons in the lobula complex (for review, see e.g. Ref. ^13^). In hoverflies and dragonflies these small target motion detector (STMD) neurons are defined by their sharp size selectivity^11, 12^. STMDs constitute a diverse group of neurons, ranging from neurons that are largely non-directional with receptive fields that encompass most of one visual hemisphere^12, 17^, to small-field, retinotopically organized STMDs^18^. In hoverflies, the dorso-frontal receptive field location of the small-field STMDs coincides with the compound eye’s dorso-frontal bright zone^19^, which has been taken as further support for a role in target pursuit^20^.

STMDs likely synapse with target selective descending neurons, TSDN^20^, which project to motor command centers in the thoracic ganglion. Like STMDs, TSDNs are defined by their sharp size selectivity^21^, but at least in dragonflies, their receptive field size, location and directionality show large diversity^22^. In dragonflies, 8 TSDN pairs provide a population code^23^ which likely drives pursuit behavior. TSDNs are particularly interesting to study as they play an important role in sensorimotor transformation. Indeed, while there are about 200,000 neurons in the *Drosophila* brain^24^, and about 20,000 neurons in the thoracic ganglion^25, 26^, only 1,000 descending neurons connect the two^27, 28^. Descending neurons are thus not only an important bottleneck in sensorimotor information transformation, but they also provide an accessible system for investigating the encoding of behaviorally relevant visual information.

In *Eristalis tenax* hoverflies, TSDNs have similar sharp size tuning^29^ to their presumed pre-synaptic counterparts^18^. The example receptive fields that have been published are located in the dorso-frontal visual field^30^, similar to the location of the bright zone^19^ and the small-field STMD receptive fields^18^. However, it is unclear how hoverfly TSDNs respond to the more dynamic stimuli experienced during pursuit. Indeed, in the field males chase targets that rapidly change flight direction and speed and they do this from a range of distances^31, 32^. In indoor experiments, males initiate pursuit of beads across a large range of physical sizes, speeds and distances, and they continue to pursue beads that briefly pause then change velocity^33^. Furthermore, in this indoor arena the male hoverflies initiate pursuit from above as well as below the bead^33^.

To understand how well matched TSDNs are to naturalistic, dynamically changing target images, we used extracellular electrophysiology of TSDNs in male *Eristalis tenax* hoverflies. We found that TSDNs (N = 118) have receptive fields in the dorso-frontal visual field, with a preferred direction up and away from the visual midline. We reconstructed the visual flow-fields as seen from the hoverflies’ position during 93 different indoor pursuits of artificial targets^33^ and recorded TSDN responses to six of these. We found that each TSDN consistently responded to the target images at unique, but specific time points. We observed a higher spike probability within the TSDN receptive field, when the target moved in the neuron’s preferred direction and had a near-optimal size. We compared the observed spike probability of the TSDN with the probability predicted from measured receptive field properties and size tuning, and found a median correlation coefficient of 0.35. However, the predicted spike probability was lower than the observed spike probability. We next compared individual TSDN responses to the population response and found that whereas individual TSDNs only responded to a small subset of each pursuit, but with high spike probability, the population of TSDNs together responded to a much larger part, but with lower population probability. This large difference between individual TSDNs could be useful if each encodes a specific behavioral outcome.

## Materials and Methods

### Electrophysiology

We recorded from 118 target selective descending neurons, TSDNs, in 112 male *Eristalis tenax* hoverflies, reared and maintained in-house as described previously^34^. At experimental time the hoverfly was immobilized ventral side up, using a beeswax and resin mixture, before an opening was made in the thoracic cavity. A small silver hook was used to elevate and support the cervical connective and a silver wire inside the opening served as a reference electrode.

Recordings were made from the cervical connective using a sharp polyimide-insulated tungsten microelectrode (2 MOhm, Microprobes, Gaithersburg, USA). Signals were amplified at 1000x gain and filtered through a 10 – 3000 Hz bandwidth filter using a DAM50 differential amplifier (World Precision Instruments, Sarasota, USA), with 50 Hz noise removed with a HumBug (Quest Scientific, North Vancouver, Canada). The data were digitized either via a 16-bit data acquisition card (NI USB6210, National Instruments, Austin, TX, USA) and the data acquisition toolbox in Matlab (Mathworks, Natick, MA, US) or a Powerlab 4/30 and LabChart 7 Pro software (ADInstruments, Sydney, Australia). In both cases the data were sampled at 40 kHz. Single units were discriminated by amplitude and half-width using Spike Histogram software (ADInstruments).

*Eristalis* males were placed ventral side up, perpendicular to, and 6.5 cm from the middle of a linearized liquid-crystal display (LCD; either PG279, Asus, or M27Q, Giga-byte Technology, both Taipei, Taiwan) with a mean illuminance of 200 Lux. The screen had a refresh rate of 165 Hz and a spatial resolution of 2,560 x 1,440 pixels (59.5 x 33.5 cm), giving a projected screen size of 155 x 138°. Visual stimuli were displayed using custom software written in Matlab (R2019b, Mathworks) using the Psychophysics toolbox^35, 36^.

### TSDN characterization

TSDNs were initially identified based on their response to a small, black, target moving over a white background^30^ (Fig. 1A). We mapped the receptive field of each neuron by scanning a target horizontally and vertically at 20 evenly spaced elevations and azimuths (Fig. 1B). The 15 x15 pixel (1.9 x 1.9°) black, square target moved at 900 pixels/s (55°/s). As the targets described in this section were not perspective corrected, the degrees correspond to the average angular size in an average receptive field (30° width, Fig. 1D), and the speed to the average angular speed across the width of the screen. There was a minimum 1 s interval between each stimulation.

**Figure 1.**
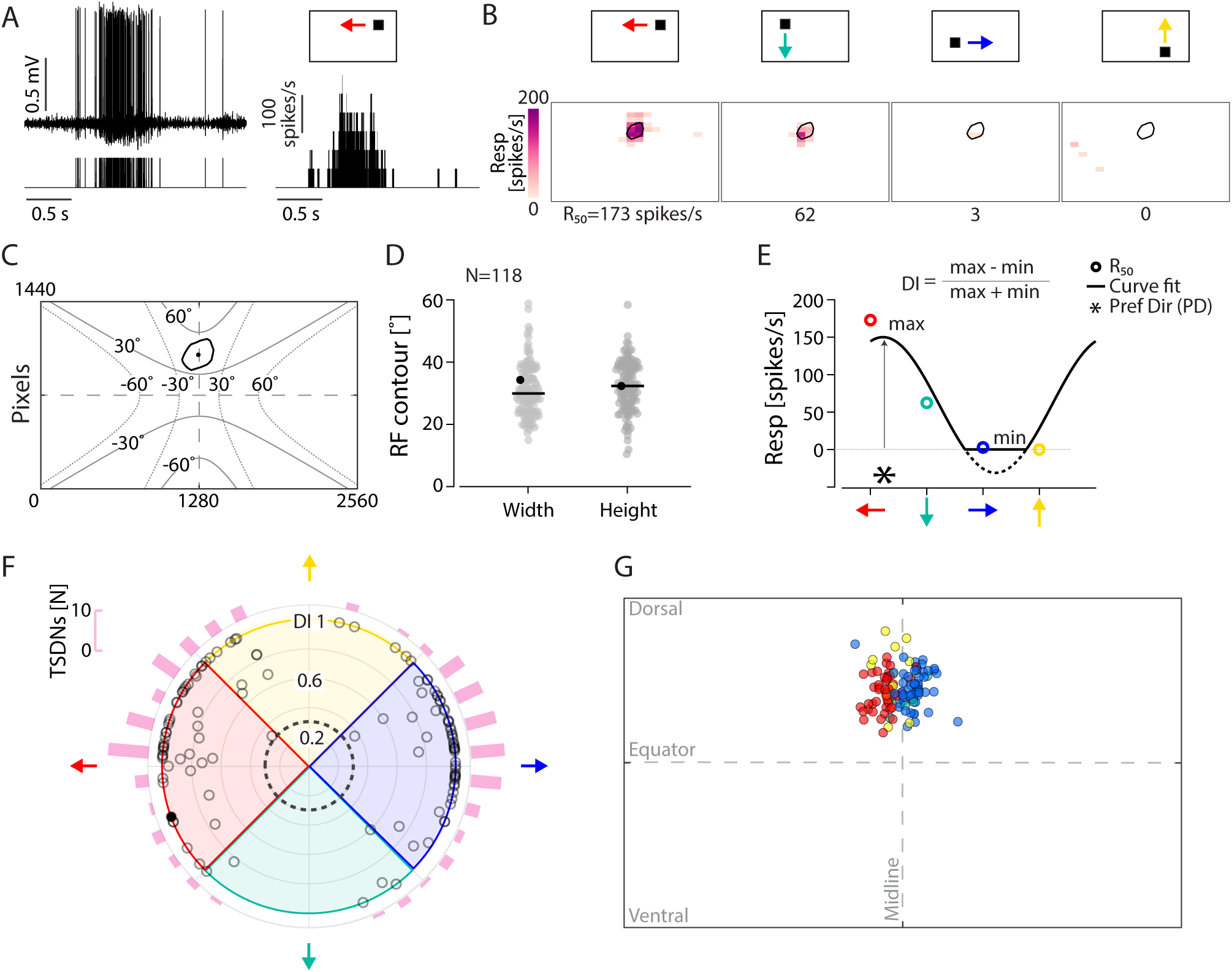
TSDNs have receptive fields in the dorso-frontal visual field and prefer motion away from the visual midline. A) Extracellular response of a single target selective descending neuron (TSDN) in a male hoverfly to a 15 x 15 pixel (1.9 x 1.9°) target scanning the screen to the left. Top left shows an example raw data trace, with the resulting spike train below, and the average spike histogram in 20 ms bins on the right. B) The response of the same example neuron to a target scanning the screen in four directions along 20 evenly spaced trajectories each. The black line indicates the 50% response contour, and R_50_ the mean response within this contour. C) The 50% receptive field response contour of the example neuron (black outline) and its receptive field center (black dot). The grey outlines illustrate the visual field in horizontal (solid lines) and vertical space (dotted lines). D) Receptive field width and height across TSDNs (N = 118), with the horizontal black line giving the median. The data from the example neuron in black. E) R_50_ to the four directions of motion (color coded from panel B), and the sinusoidal curve fit (black line). The preferred direction (Pref Dir, PD, *) and directionality index (DI) as obtained from the curve fit. F) The data from 118 TSDNs, where the distance from the center indicates directionally index, and the location along the circumference indicates preferred direction. The data from the example neuron in black. The dashed circle shows a directionality index of 0.3, used to separate directional neurons from non-directional^18^. The pink histogram along the circumference shows the preferred direction across TSDNs (10° bins), with 44 neurons preferring leftward motion, 52 rightward, 16 upward, and 6 downward motion. This distribution was significantly non-uniform (*P* < 0.0001, Rayleigh test). G) The receptive field center locations color coded according to their preferred direction (N = 118).

**Figure 2.**
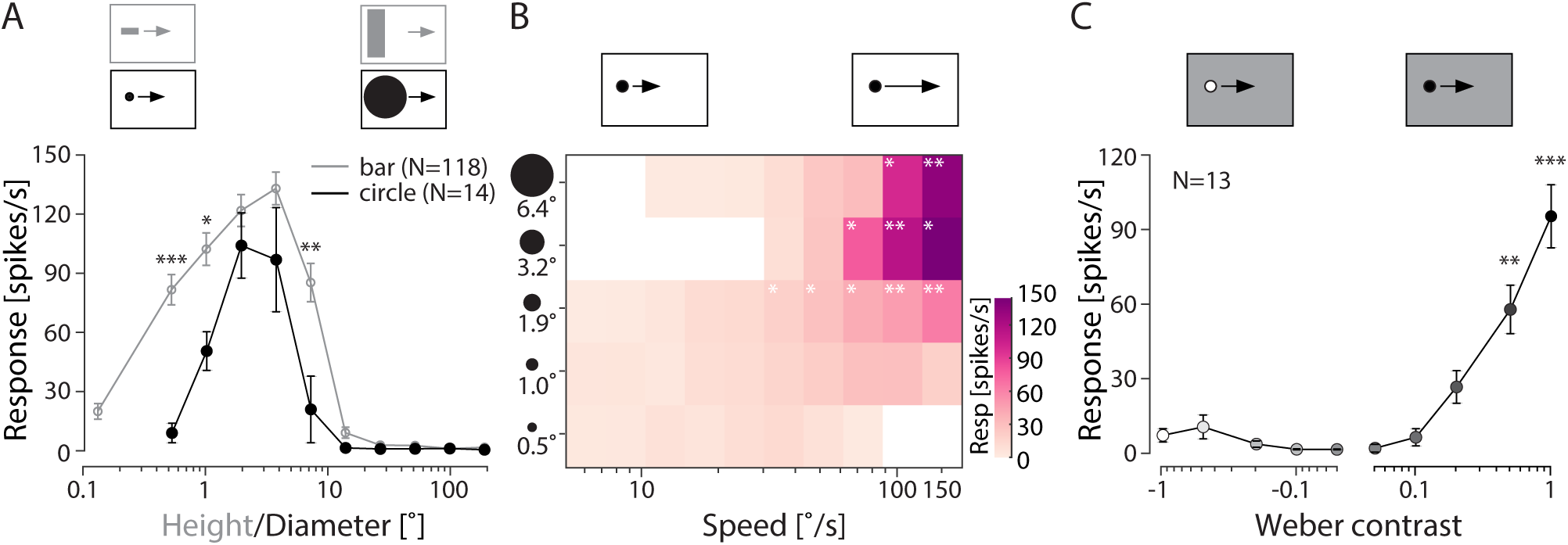
TSDNs respond to small, dark contrast, moving targets. A) The response to targets scanning the screen at 900 pixels/s (55°/s) horizontally through each neuron’s receptive field. The grey data show responses to different bar heights, where the width was fixed at 15 pixels (1.9°, N = 118). The black data show the responses to circles with different diameters (N = 14). The data are displayed as mean ± sem. Significance is shown using two-way ANOVA followed by Šídák’s multiple comparisons test with \**P* < 0.05, \*\**P* < 0.01 and, \*\*\**P* < 0.001. B) The response to circular, black targets scanning the screen horizontally through each neuron’s receptive field. The x-axis indicates the speed, the y-axis the diameter of the target, and the response is color coded to the mean response across neurons (N = 9, 7, 14, 7, and 9 for sizes of 0.5° to 6.4°, respectively). Significance was investigated using two-way ANOVA followed by Dunnett’s multiple comparisons test, \**P* < 0.05 and \*\**P* < 0.01. C) The response to circular targets scanning the screen horizontally over a uniformly grey background. The data are displayed as mean ± sem (N = 13). Significance is shown using one-way ANOVA followed by Dunn’s multiple comparisons test with \*\**P* < 0.01 and \*\**P* < 0.001.

For each trajectory we calculated the mean spiking frequency within 20 bins (142 ms each), assuming a neural response delay of 18 ms. This allowed us to create a 20 x 20 grid of the response to each direction of motion (Fig. 1B). We then calculated the average response to the four directions of motion and interpolated this to a 100 x 100 grid. We used Matlab’s *contour* function to find the 50% response, and quantified receptive field height and width. We defined the center of the receptive field as the center of the 50% contour line using Matlab’s *centroid* function.

We calculated the average spike frequency to each direction of motion within the 50% contour line (*R*_50_, Fig. 1B) and fit a sinusoidal function to the response to extract its preferred direction (PD, Fig. 1E). We calculated a directionality index (DI, Fig. 1E) for each neuron based on the maximum and minimum of the curve fit. If the minimum was below 0, we set it to 0, as neurons cannot have a negative spike frequency.

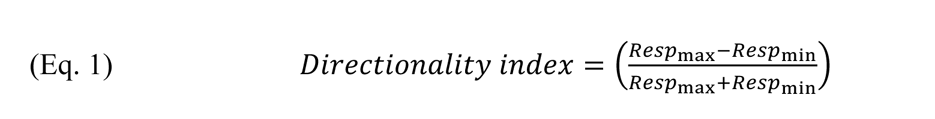

We performed k-means clustering on the observed receptive fields using the *evalclusters* function in Matlab. For this, we used the receptive field center location along the horizontal and vertical axes, the receptive field width, height, preferred direction, and directionality index. We estimated the optimal cluster number under five conditions: 1) when using all these receptive field parameters, 2) when using only the horizontal receptive field center location and the preferred direction, 3) using all parameters except the horizontal receptive field center and the preferred direction, 4) using all parameters except the horizontal receptive field center, and 5) using all parameters except the preferred direction.

To determine size selectivity, we drifted a black bar over a white background at 900 pixels/s (average 55°/s) in each neuron’s horizontal preferred direction (i.e. left or right) through the center of its receptive field. The bar side parallel to the direction of travel was fixed at 15 pixels (average 1.9°) whilst the perpendicular side varied from 0.12° (1 pixel) to the full height of the screen. In a subset of neurons, we used a black circle drifting horizontally at 900 pixels/s. The circle diameter varied from 0.51° (4 pixels) to the full height of the screen. There was a minimum 4 s interval between each stimulation.

To determine the speed sensitivity function, a black circle drifted over a white background in each neuron’s preferred horizontal direction, through the center of its receptive field. We used circle diameters of 4, 8, 15, 25, and 50 pixels, corresponding to an average 0.5°, 1.0°, 1.9°, 3.2° and 6.4°, which represent typical target sizes at the start of pursuits ^33^. The speeds were spaced logarithmically from 100 to 2475 pixels/s (average 6°/s to 150°/s). There was a minimum 4.4 s interval between each stimulation.

To determine the contrast sensitivity function, a circle with a diameter of 15 pixels (1.9°) moved at 900 pixels/s horizontally through the center of each neuron’s receptive field, across a mid-luminance, grey background. The Weber contrast varied from -1 to 1, using the following equation:

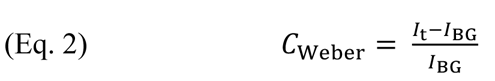

where *I*_t_ is the intensity of the target, and *I*_BG_ the intensity of the background.

For size, speed and contrast sensitivity functions, we quantified the average spike frequency within a 142 ms analysis window, to match the receptive field analysis, centered on each neuron’s receptive field. We repeated the speed and contrast sensitivity experiments at least three times in each neuron. When stimuli were repeated several times, we calculated the mean from each neuron. All experiments were presented in random order.

### Reconstruction of retinal flow fields from target pursuits

We used behavioral pursuits of artificial targets (black beads, 0.6, 0.8, 1 and 3.85 cm in diameter) filmed with two cameras at 120 frames/s, followed by 3D reconstructions of the hoverfly and target trajectories over time (Fig. 3A, for details see Ref. ^33^). In the 3D position data, the x-and y-axes define the two sides, and the z-axis the height (Fig. 3A). *F* is used to describe the hoverfly and *B* the bead.

**Figure 3.**
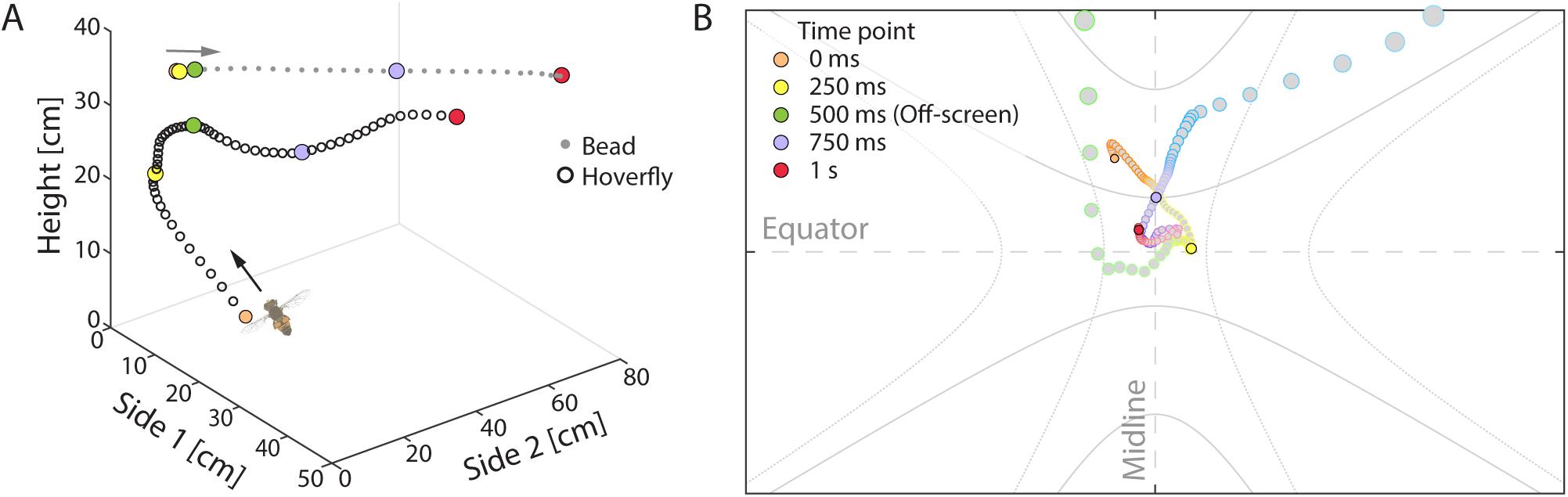
Reconstructing the retinal image of target pursuits. A) An example extract of a male *E. tenax* (circles) pursuing an 0.8 cm diameter bead (dots). The data show the locations of the hoverfly and bead at 60 Hz resolution, color-coded from orange to red to indicate the time every 250 ms. B) The retinal image of the target from the same example trajectory, as displayed at 165 Hz resolution on our 2D visual screen, for use in electrophysiology. The reconstruction is color coded to indicate time.

**Figure 4.**
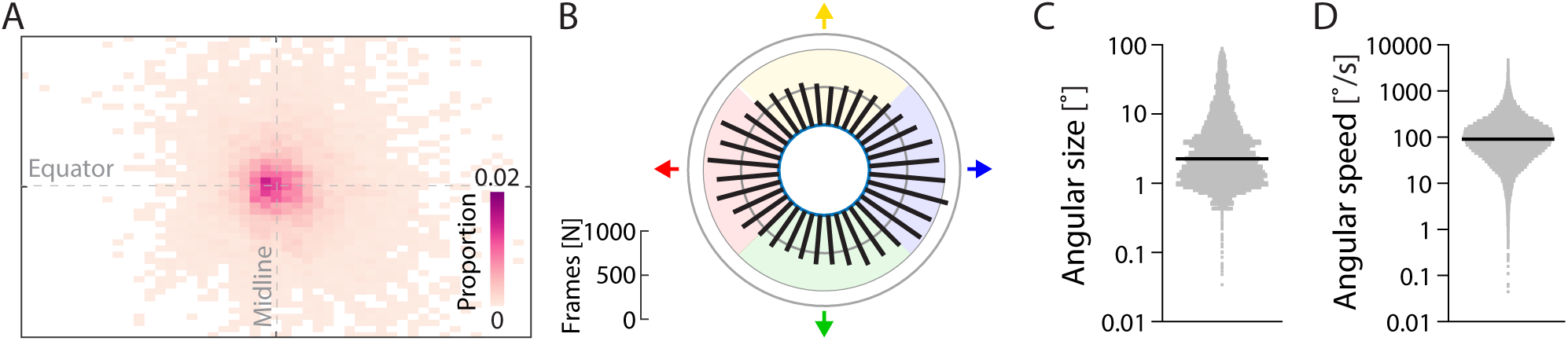
The retinal image from 93 target pursuits as displayed on the 2D visual screen. A) The probability that the target image appeared in each grid unit (50 x 50 resolution) on our visual screen. B) The direction of target motion, in 10° bins. The distribution was significantly non-uniform (*P* < 0.0001, Rayleigh test). C) The distribution of the targets’ instantaneous angular size from all 93 pursuits interpolated to 165 Hz, from Ref. ^33^. D) The distribution of the targets’ instantaneous angular speed from all 93 pursuits interpolated to 165 Hz, from Ref. ^33^.

To reconstruct the retinal flow fields to be projected onto our 2D screen (Fig. 3B), we needed to adjust for the hoverfly’s immobilized position in front of the screen. We therefore situated the hoverfly in the origin point in a 3D space (0, 0, 0) using subtraction at each point in time (Fig. S2A). We next determined the relative angle (α) between the vector describing the hoverfly heading (*H*_flight_) and a fixed heading unit vector (*H*_fixed_, (0, 0, -1), Fig. S2B). The relative angle (α) was calculated using the law of cosine:

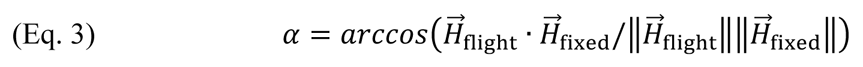

where ‘·’ is the dot product and ‘|| ||’ is the Euclidean norm. We took the relative angle (α) to produce a rotation matrix using Rodrigues’ formula (Fig. S2C):

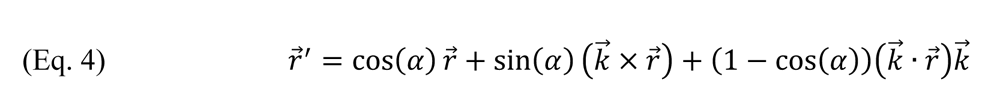

where ‘×’ is the cross product, *ř* is the vector to the original bead position, *ř*′ is the vector to the new bead position (*x’*_B_*, y’*_B_*, z’*_B_) and *K*^7⃗^ is the unit vector in the direction of the rotation axis:

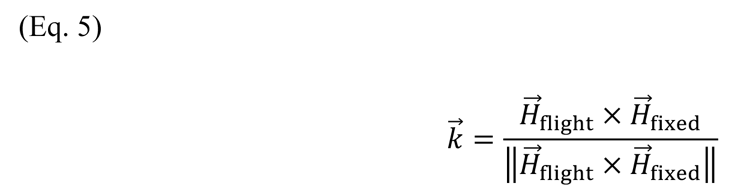

The data sequence was smoothed using Matlab’s *lowess* smoothing, which is a locally weighted linear regression method, using 2% of the total number of data points. The data were resampled from 120 Hz to 165 Hz to match the refresh rate of the screen, using Matlab’s *resample* function.

To display the reconstructed target on the 2D screen for electrophysiology, where *H*_fixed_ was centered on the display (Fig. S2D), we first determined where the target would be projected.

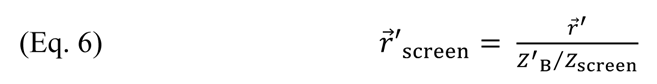

where *z*_56*((7_ = 6.5 cm, which is the distance between the fixed hoverfly and the center of the screen.

To calculate the perspective corrected diameter of the target on the 2D screen (*w*_screen_) we used the distance between the immobilized hoverfly and the bead’s location on the screen (‖*ř*^4^_56*((7_‖, Fig. S2E), the length of the vector *ř*′ (Eq. 4), and the physical size of the bead (*w*, 0.6, 0.8, 1, or 3.85 cm diameter):

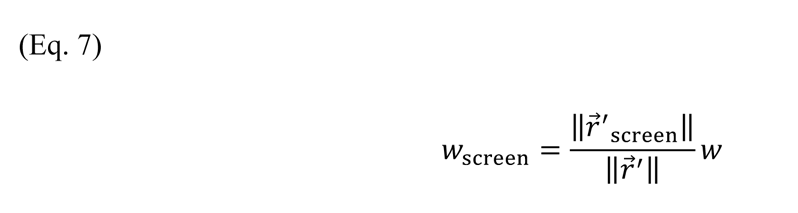

For electrophysiology the resulting 2D stimuli were displayed as black circles on a white background.

### The retinal image of the target

To determine the probability of the reconstructed bead image to be displayed in different locations on the 2D visual screen, we used the target location (Eq. 6) from all frames of all 93 pursuits. We divided our visual screen into a 50 x 50 grid and calculated the proportion of occurrences within each grid unit.

We quantified the direction the reconstructed bead image moved between each frame of the 93 pursuits using the positions on the screen (*ř*′_56*((7_, Eq. 6). The direction was determined between two consecutive frames, using the law of tangents.

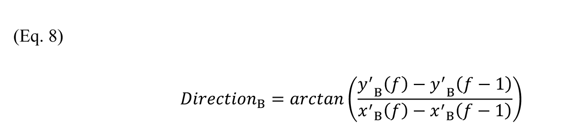

where *f* is each frame number.

We replotted the angular size and speed across the 93 pursuits from Ref ^33^ after interpolating the data to 165 Hz.

### Observed spike probability

For electrophysiology, we used six of the 93 pursuits, where each pursuit started either below (Fig. 6FG *i*-*iii*) or above (Fig. 6FG *iv*-*vi*) the 0.8 cm diameter bead, and lasted for at least 2.9 s. Each pursuit was shown at least nine times to each neuron, in a pseudo-random order, with a minimum 2 s rest between presentations. From the repetitions we quantified the observed spike probability, defined as the fraction of trials eliciting at least one spike within a given stimulus frame, where each frame was 6.1 ms long, as the screen refresh rate was 165 Hz. In this analysis, we assumed an 18 ms delay between stimulus presentation and neural response, i.e. 3 frames.

For each TSDN, and each trajectory, we calculated the percent of target images that generated an observed spike probability above 0.5. For the population of TSDNs (N = 27), we calculated the percent of target images that generated an observed spike probability above 0.5, in any neuron, for each trajectory (Fig. S6A-C).

To calculate the population spike probability, we quantified the fraction of TSDNs that gave an observed spike probability above 0.5, for each frame, for each trajectory (Fig. S6D-F).

### Predicted spike probability

We determined a predicted spike probability for each frame of the reconstructions based on each neuron’s receptive field and size tuning, together with the target’s location on the screen, its direction of travel, and its size. To do this practically, we first used the receptive fields recorded from each neuron in response to the four directions of motion (Fig. 1B) to estimate the receptive field shape for all directions of target motion (at 1° resolution), assuming sinusoidal direction tuning (see e.g. Fig. 6A), and normalized these to a maximum of 1 (Fig. 6B). For each target location and direction we could now extract a normalized response (*R*_norm_, Fig. 6C). Finally, by incorporating the size tuning for each neuron (*R*_size_, Fig. 6D), we can estimate a spike rate, *λ* (Fig. 6E), which incorporates the target’s location, its direction, and size, using multiplication:

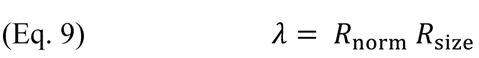

**Figure 5.**
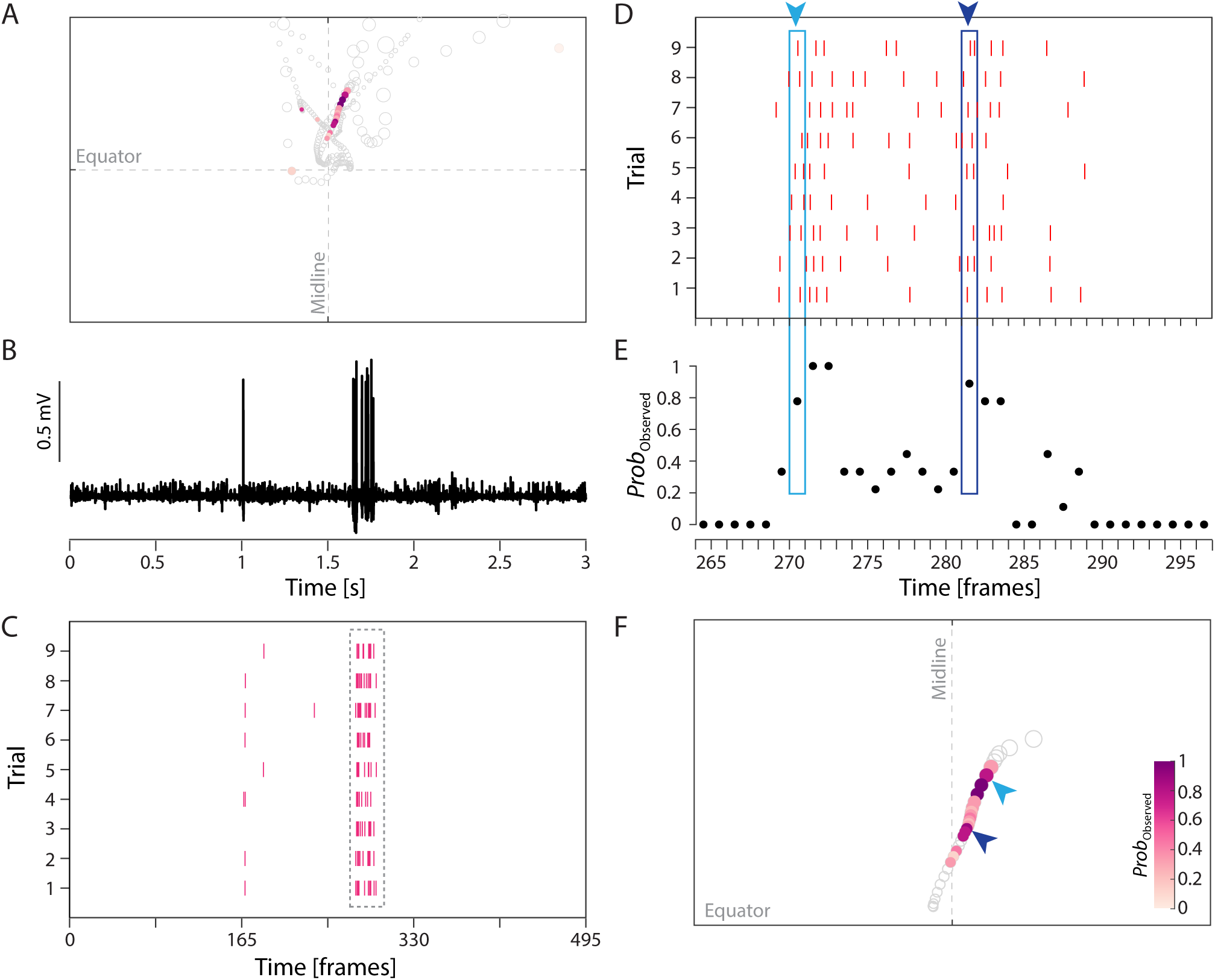
Observed spike probability quantification. A) An example reconstructed target pursuit (same as in Fig. 3) with grey circles showing the location of the target in each frame (165 Hz resolution). For clarity, they have here been scaled to twice their size. The color coding indicates an example TSDN neuron’s observed spike probability (color bar in panel F). B) An example extracellular recording from a TSDN responding to the reconstructed pursuit. C) A raster plot for the example neuron to nine trials of the same reconstruction. The data in panel B are shown in trial 1. D) A magnification of the dashed area from panel C. For each frame, we counted the number of trials that generated at least one spike. In this example, during frame 270 (pale blue), 7 of 9 trials generated at least one spike, and during frame 281 (dark blue), 8 of 9 trials generated a response. E) The observed spike probability, defined as the fraction of trials that generated at least one spike. F) Visualization of observed spike probability (color coded) for the target pursuit reconstruction in panels D and E.

**Figure 6.**
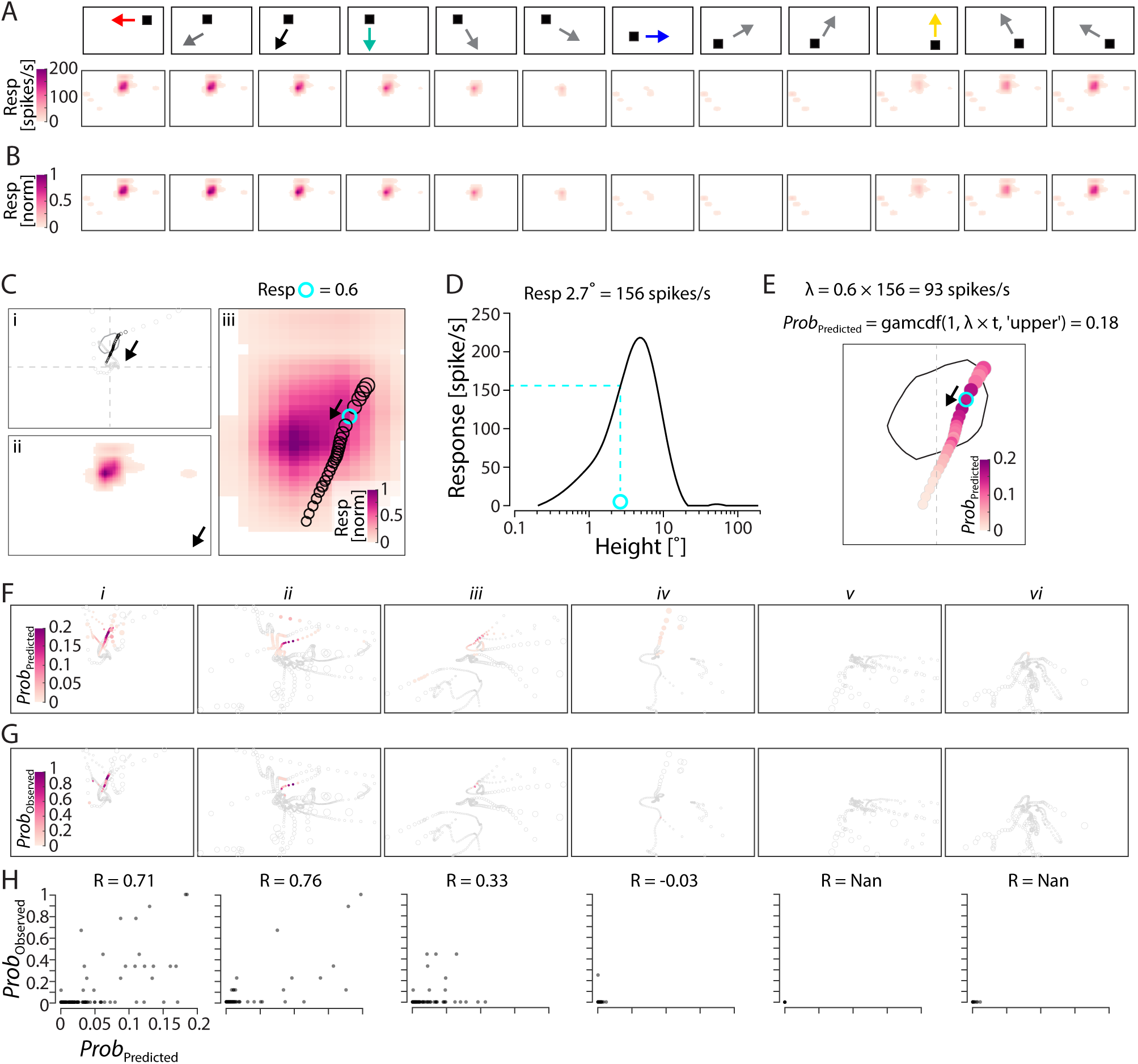
Calculation of predicted spike probability. A) We simulated the receptive field for all possible target directions (at 1° resolution) assuming sinusoidal direction tuning, here shown at 15° resolution. B) The normalized receptive fields. C) *i*) Extract of an example target trajectory, and a neuron’s 50% receptive field response contour. *ii*) The normalized receptive field for the direction of target motion shown in panel *i. iii*) Panels *i* and *ii* magnified and superimposed. The normalized response for the cyan example frame is 0.6. D) The expected spike frequency for a 2.7° target (the diameter of the cyan example), 156 spikes/s, extracted from this neuron’s size tuning. E) The predicted spike frequency, 11, used to predict that there was a 0.18 chance that this cyan frame would generate at least one spike. The pictogram shows the predicted spike probability for a subset of the reconstructed pursuit (color coded). F) Predicted spike probability for the same TSDN to six different reconstructed target pursuits. G) The observed spike probability for the same TSDN to the same six target pursuits. H) Correlation between the observed and predicted spike probability for each trajectory. Each dot shows the result for an individual frame.

From *11* we calculated the predicted spike probability, which is the probability that at least one spike would occur within a given frame. For this purpose, we first examined the spike-time statistics by calculating the inter-spike interval (ISI) in each TSDN’s response to a target scanning the screen in the preferred horizontal direction (as in Fig. 1A, B). We used Matlab’s *fitdist* to fit the ISI distribution with either a gamma or a Poisson function, as these two are often used to described spike-time statistics^37, 38^. We then used Matlab’s *gfit* to calculate the root mean squared error (RMSE) between the fit and the ISI data and found that the gamma function provided a better fit than the Poisson function (Fig. S3). To calculate the predicted spike probability, we therefore used Matlab’s *gamcdf* function:

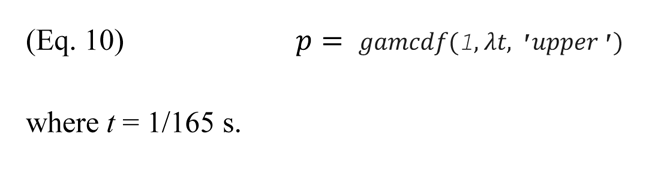

### Data analysis and statistics

All neurons that were sharply tuned to small, black targets (Fig. 2A), and did not respond to a looming stimulus^30^, were kept for further analysis. All data analysis was performed in Matlab (R2020b, Mathworks) and Prism 7.0c for Mac OS X (GraphPad Software).

Correlation between predicted and observed spike probability was calculated for each neuron using Matlab’s *corrcoef* either by treating each trajectory data separately (*i-vi*, Fig. 7) or after combining all data for the six trajectories (All, Fig. 7). For statistical comparison between the percent of target images that generated an observed spike probability above 0.5 in either individual TSDNs or the population (Fig. 8A), we used a bootstrapping method. For this, we randomly selected 20 TSDNs, quantified the percent of target images that generated an observed spike probability above 0.5 in any neuron, and repeated this process 27 times. To reduce the differences in power when comparing non-zero observed spike probabilities in individual TSDNs (N = 3,629) with non-zero population probabilities (N = 159, Fig. 9B) we randomly selected 159 observed spike probability occurrences before statistical analysis. For all analyses, the sample size, type of test and *P*-values are indicated in each figure legend.

**Figure 7.**
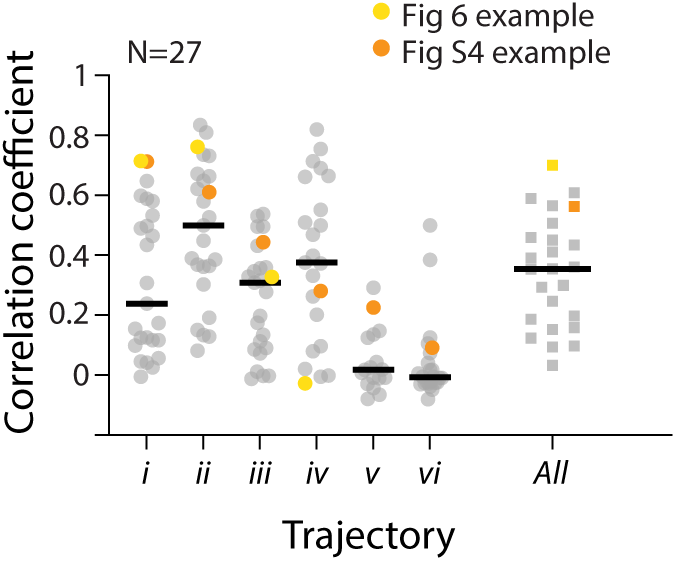
The predicted spike probability is correlated with observed spike probability. The correlation coefficients between observed and predicted spike probability of TSDN neurons (N = 27) for the six trajectories (*i-vi*) or when analyzed together (All). The median correlation coefficients were 0.24, 0.5, 0.31, 0.38, 0.02, -0.01 and 0.35. Color coding shows example neurons from Figure 6 (yellow) and Figure S4 (orange).

**Figure 8.**
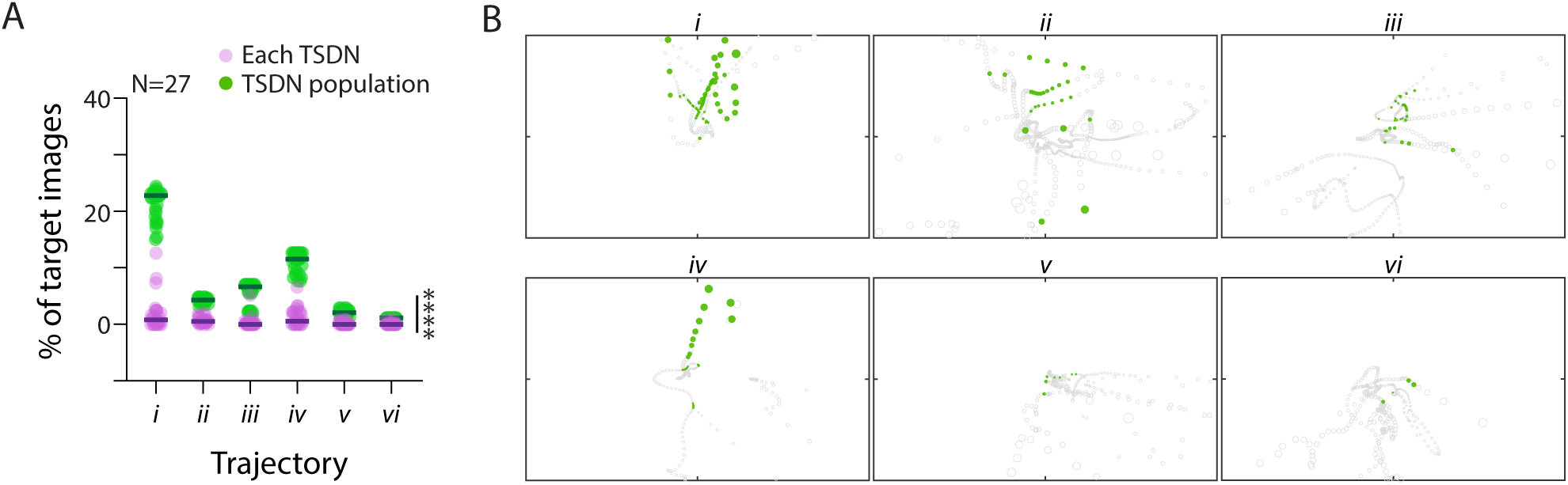
The population of TSDNs respond to a larger part of the pursuit. A) The percent of frames that generated above 0.5 observed spike probability, in each individual neuron (purple), and in the population of neurons (green), for each of the six trajectories (*i-vi*). The population data were acquired by randomly selecting 20 neurons 27 times (see Methods). Significance was investigated using the Kolmogorov-Smirnov test: \*\*\*\**P* < 0.0001. B) Target images which had above 0.5 observed spike probability in at least one TSDN (N = 27) are color coded in green.

**Figure 9.**
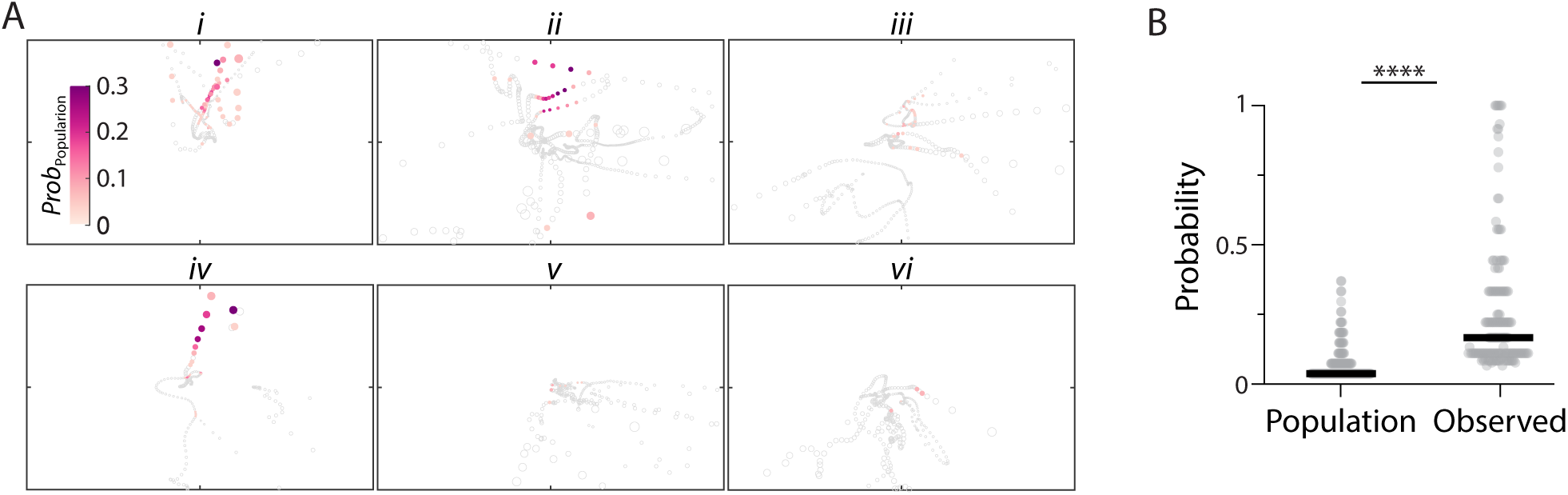
TSDN observed spike probability is higher than the population spike probability. A) The population spike probability color coded for each of the six trajectories (*i-vi*). B) The population spike probability (median 0.04) was significantly lower than the observed spike probability (median 0.17). Significance was investigated using the Kolmogorov-Smirnov test: \*\*\*\**P* < 0.0001.

### Data and code availability

All data and analysis scripts will be submitted to DataDryad.

## Results

### TSDN receptive fields suggest a homogenous population

Target selective descending neurons (TSDNs) in male *Eristalis tenax* hoverflies often have no spontaneous activity^30^ but respond vigorously to the motion of small dark targets (example shown in Fig. 1A). The receptive field of an example TSDN shows strong responses to leftward and downward motion in the dorso-frontal visual field (Fig. 1B). For each neuron, we calculated the average receptive field, determined the 50% response contour (same example neuron, black outline, Fig. 1C), and quantified receptive field width and height.

Across 118 TSDNs we found that the receptive field widths ranged from 15° to 59° (minimum to maximum), with a median of 30° (Fig. 1D). The height varied from 10° to 58°, with a median of 32° (Fig. 1D). The TSDN receptive fields are thus substantially larger than the small-field STMDs, which have halfwidths of 7-8° ^18^.

We next obtained the mean spike rate within the 50% response contour (R_50_, Fig. 1B) to extract each neuron’s preferred direction (Pref Dir, PD, Fig. 1E) and directionality index (DI, Fig. 1E). For comparison across neurons, we display each neuron’s preferred direction in a polar plot, where the distance from the center is correlated with the directionality index (open symbols, Fig. 1F). The dashed line shows the cut-off previously used to separate directional from non-directional STMDs^18^. Using this definition, all TSDNs are directional, with directionality indices ranging from 0.33 to 1, and a median of 1 (Fig. 1F). The pink histograms along the circumference show the distribution of preferred directions, which was significantly non-uniform (*P* < 0.0001, Rayleigh test).

To determine if the preferred direction was correlated with location in the visual field, we plotted the receptive field centers (black dot, Fig. 1C) for all 118 TSDNs, color coded according to their preferred direction (Fig. 1G). We used clustering of the 118 TSDNs’ receptive field center’s horizontal location, vertical location, the receptive field width, height, preferred direction and directionality index, and found that the optimal cluster number was two (“All”, Fig. S1A). We next repeated the clustering, but only included the receptive field center’s horizontal location (X) and preferred direction (PD), and again found that the optimal cluster number was two (“X + PD”, Fig. S1A). If we excluded the receptive field center’s horizontal location (X) and/or the preferred direction (PD), the optimal cluster number was larger (Fig. S1A). We thus conclude that the 118 TSDNs could be divided into two clusters based on preferred direction and the receptive field horizontal location. The two clusters are placed on one side each of the visual midline in the dorso-frontal visual field (Fig. 1G, S1B), suggesting that they constitute the right-and left-hand version of the same population. Their preferred direction of motion is away from the visual midline (Fig. 1G).

### TSDNs are tuned to the dark contrast, small targets

TSDNs, like STMDs, are defined by their sharp size selectivity (grey data, Fig. 2A), with no response to targets subtending more than 10° ^20^. We compared the response to bars and circles and found that the response to bars peaked at 3.6° height (grey data, Fig. 2A), and that the peak response to circles was at 1.9° diameter (black data, Fig. 2A). The response to bars was significantly larger than the response to circles of 0.5°, 1° and 7° diameter (two-way ANOVA, *P* = 0.0006, *P* = 0.046, *P* = 0.0035, respectively). The lower response to the 7° circle could be accounted for by recruitment of more lateral inhibition^39, 40^. The lower response to the 0.5° and 1° circles could be because they had a smaller area, and thus generated lower neural contrast^12^.

We next investigated the speed response function to circles of different diameters, as previous STMD work has shown a response shift to higher velocities when using larger objects^17^. We found that the response depended significantly on both diameter and speed (two-way ANOVA, diameter: *P* = 0.0147; speed: *P* < 0.0001; diameter x speed: *P* < 0.0001, Fig. 2B). We found that the response was significantly larger than the spontaneous rate (stars, Dunnett’s multiple comparisons test, Fig. 2B), for the three larger bead sizes at the higher speeds, and that significance shifted to higher speeds with bead size (stars, Fig. 2B).

We investigated the contrast response function, as dragonfly and hoverfly STMDs have been shown to prefer dark contrast targets^12, 41^, and found that the response depended significantly on contrast and was significantly higher than the spontaneous rate at 0.5 and 1 Weber contrast (Fig. 2C, N = 13, one-way ANOVA, *P* < 0.01 and *P* < 0.001). When the target was brighter than the background, the response was negligible.

### Reconstructing the retinal image from pursuits of artificial targets

We next reconstructed visual flow fields as experienced by male hoverflies during pursuits of artificial targets in an indoor arena^33^. We used 93 pursuits filmed with two cameras at 120 frames/s, followed by 3D reconstructions of the hoverfly and the bead positions over time^33^. The example data show a subset of a pursuit of an 0.8 cm diameter black bead (Fig. 3A).

To reconstruct the visual stimulus as observed from the position of the immobilized hoverfly, we first calculated the bead position relative to the hoverfly heading using Rodrigues’ formula (Fig. S2A - C). We interpolated this at 165 Hz, the refresh rate of our screen. We then calculated each target position — as it needed to be projected onto the 2D screen — and its perspective corrected size in pixels (Fig. S2D, E), to complete the reconstruction of the retinal flow field (example trajectory, Fig. 3B).

Across all frames of all 93 pursuits^33^ we found that the target image was most frequently projected in the center of the screen (Fig. 4A). We found that the bead moved in all directions on the visual screen, but that the distribution was significantly non-uniform with a preference for horizontal motion (Fig. 4B, *P* < 0.0001, Rayleigh test). For comparison with neural data, we replotted the angular size of the target image from Thyselius et al.^33^ and found that it ranged from 0.03° to 90°, with a median of 2.2° (Fig. 4C), matching the TSDN size tuning (Fig. 2A). The angular speed ranged from 0 to 4800°/s, with a median of 90°/s (Fig. 4D), where we expect the TSDNs to respond (Fig. 2B).

### TSDN responses to reconstructed target pursuits

The data above show that TSDN receptive fields (Fig. 1), size (Fig. 2A) and speed response functions (Fig. 2B) overlap with the target images from pursuits as projected onto the visual screen (Fig. 4). We next asked, how do TSDNs respond to reconstructed pursuits? For this purpose, we chose six of the 93 reconstructions (one example shown in Fig. 3, 5A). The raw data (Fig. 5B) shows an example TSDN response to this trajectory.

We recorded the response to at least 9 trials in each TSDN (Fig. 5C) to calculate an observed spike probability, defined as the fraction of trials that generated at least one spike within each frame (6.1 ms time window, as the screen refresh rate was 165 Hz, Fig. 5D, E). For example, during frame 270, this TSDN responded with at least one spike in seven of the nine trials (Fig. 5D, light blue), giving an observed spike probability of 0.77 (Fig. 5E, light blue). During frame 281, the observed spike probability was 0.88 (dark blue, Fig. 5D, E). To visualize the observed spike probability, we color-coded the reconstructed target pursuit (Fig. 5A; zoomed in, Fig. 5F).

We found that this TSDN only responded to a small subset of frames in the pursuit (Fig. 5A, F). Could this response be predicted from the receptive field, directionality and size tuning (Fig. 1, 2)? To answer this question, we used this neuron’s interpolated receptive field (every 30° shown in Fig. 6A, B), and size tuning (Fig. 6D), to predict a spike rate (for details, see Methods). In the example pursuit extract shown in Figure 6C, the target moves down and slightly to the left, giving a normalized response of 0.6 for the cyan frame. As this cyan example had a 2.7° diameter, we extract a spike rate of 156 spikes/s (Fig. 6D). These two values are multiplied to get the predicted spike rate, λ (93 spikes/s, Fig. 6E), giving a predicted spike probability of 0.18 (Fig. 6E).

Across the pursuit, the example TSDN is predicted to respond to a small subset of target images in the first three trajectories, but not to the last three, probably because they are predominantly projected in the ventral visual field (Fig. 6F, and see Fig. S4A for second example TSDN). A qualitative comparison suggests that this is quite similar to the observed spike probability (Fig. 6G, S4B). We quantified this qualitative comparison by determining the correlation coefficient between the observed and the predicted spike probability for each pursuit (example neurons in Fig. 6H, S4C). This showed correlation coefficients up to 0.76, depending on trajectory (Fig. 6H). Across 27 TSDNs we found that the median correlation coefficients ranged from -0.01 to 0.5, depending on trajectory (Fig. 7). Analyzed together, the median correlation coefficient was 0.35 with lower and upper confidence limits of 0.2 and 0.45 (“All”, Fig. 7).

We noted that even if the observed spike probability could reach 1, the largest predicted spike probability was often lower (see e.g. Fig. 6H, S4C). Across 27 TSDNs the maximum observed spike probability ranged from 0.44 to 1, with a median of 1. This was significantly larger than the maximum predicted spike probability, which ranged from 0.04 to 1, with a median of 0.46 (Fig. S5, *P* < 0.0001, Wilcoxon test). Taken together this suggests that TSDN directionality, size tuning and receptive field location can predict some of the response to reconstructed target pursuits.

### Response to limited part of the pursuit

As mentioned above, only a limited part of the reconstructed trajectories generated a response (Fig. 5A, 6G, S4B). To quantify this, as a conservative estimate we determined the percentage of frames that generated an observed spike probability above 0.5 in each TSDN (Fig. S6A). We found that within each of the 27 TSDNs, the percentage of frames that generated an observed spike probability above 0.5 varied between 0 and 12.6%, with medians of 0.8, 0.5, 0, 0.5, 0, and 0% in the six trajectories (purple, Fig. 8A).

We also noted that individual neurons responded to different parts of the pursuit (compare e.g. Fig. 6G with Fig. S4B). We therefore determined the percentage of frames that had an observed spike probability above 0.5 in at least one neuron (Fig. 8B, S6B-C, and see methods). This analysis shows that the TSDN population responded to a substantially larger part of the reconstructed pursuits. Indeed, the medians were 23, 4.3, 6.6, 11.5, 2.1, and 1.1% for the six different trajectories (green, Fig. 8A). This was significantly larger than the percentage of frames that generated a response in each individual neuron (compare green and purple, Fig. 8A, *P* < 0.0001 Kolmogorov-Smirnov test).

### Population spike probability is lower than the observed spike probability in individual TSDNs

The much larger response across the TSDN population compared with individual TSDNs (Fig. 8A) suggests that there is a large variation between neurons. To analyze this further, we determined the population spike probability, defined as the fraction of TSDNs that generated an observed spike probability above 0.5 to each frame (Fig. S6D-F). We found that the population spike probability was higher in the dorso-frontal visual field, especially in response to trajectories *i, ii*, and *vi* (Fig. 9A). The population spike probability across trajectories ranged from 0.04 to 0.37, with a median of 0.04, meaning that only one TSDN responded (Fig. 9B, N = 159 frames). A population spike probability of 1 means that every TSDN responded to that frame with at least one spike, whereas a population spike probability of 0 means that no TSDN responded. We compared this to the observed spike probability from each neuron (see Fig. 5, 6), which ranged from 0.07 to 1 with a median of 0.17 (Fig. 9B). An observed spike probability of 1 means that the individual TSDN responded with at least one spike to a specific frame to every repetition of the stimulus (Fig. 5).

As the observed spike probability was significantly larger than the population spike probability (Kolmogorov-Smirnov test, *P* < 0.0001, Fig. 9B), this means that individual neurons responded more consistently to individual frames within the pursuit, than the TSDN population as a whole did. This suggests that each TSDN responded to a different part of the pursuit. Together with the finding that the TSDN population responded to a larger part of the pursuit (Fig. 8), this suggests that the TSDNs could provide a sparse population code. Indeed, population codes have been shown in e.g. dragonfly TSDNs and salamander ganglion cells^7, 23^.

## Discussion

We here show that hoverfly TSDNs have receptive fields with a 30° half-width, with centers located in the dorso-frontal visual field (Fig. 1). The neurons are highly directional, preferring motion away from the visual midline (Fig. 1G). We reconstructed the target image (Fig. 3, 4, S2) from pursuits of artificial targets^33^, and show that each neuron’s receptive field location, directionality and size tuning (Fig. 1-2) can be used to predict the neural response (Fig. 5-7, S4). However, in all cases the predicted spike probability to individual frames within the reconstructed pursuit was lower than the observed spike probability (Fig. S5). We showed that while individual TSDNs responded to a small subset of frames (median below 1%, Fig. 8A) within the reconstructed pursuits with high spike probability (median 0.17, Fig. 9B), the TSDN population responded to a larger proportion of each reconstructed pursuit (median up to 23%, green, Fig. 8, S6A-C) but with lower population spike probability (median 0.04, Fig. 9, S6D-E).

### Comparison with STMDs

TSDNs are likely post-synaptic to the STDMs that have been described in the hoverfly and dragonfly lobula (see e.g. Ref. ^13, 20^). Hoverfly STMDs constitute a diverse group of neurons that are united by their sharp size selectivity^20^. Indeed, hoverfly STMDs include neurons that are highly directional as well as those that are non-directional, the receptive fields range from small-field to those that encompass most of one visual hemisphere, and their ability to respond to targets in clutter varies^12, 18^. However, based on receptive field properties, the TSDNs that we recorded here appear to come from only one population (right-and left-hand side, Fig. S1). We identified potential TSDNs by their strong response to small, dark-contrast targets scanning the visual monitor (Fig. 1A). It is possible that there are other TSDNs that we have missed, due to this initial selection. Nevertheless, this comparatively homogenous TSDN population (Fig. 1, S1) is intriguing, considering the diversity of the presumably pre-synaptic STMDs^12, 18^.

We quantified the speed response function to targets of different diameter, and found that the response increased with target speed, for the target sizes used here (Fig. 2B). This is different to dragonfly STMDs^17^, which showed a dramatic shift to higher speeds when using an 8° wide target (peak at 200°/s) compared with a 0.8° square target (peak at 60°/s). This therefore suggests that the speed response function differs between the lobula STDMs and the TSDNs. This is interesting as there are other notable differences between optic lobe neurons and the descending neurons. For example, widefield sensitive neurons in the optic lobe show substantial post-excitatory inhibition^42, 43^, whereas widefield sensitive descending neurons display post-excitatory persistent firing^44^. In addition, while many STMDs respond robustly to targets against velocity matched background clutter^12, 45^, TSDNs do not^29, 46^.

STMDs and TSDNs are believed to get their input from elementary STMDs (ESTMDs), which create selectivity to the motion of small, dark-contrast targets^39, 41, 47^. Indeed, the TSDN contrast response function (Fig. 2C) was similar to those recorded in dragonfly and hoverfly STMDs^12, 47^, with very little response to bright contrast targets. This is interesting as during target pursuit in the field, territorial intruders or conspecific females are often displayed as dark contrast objects against the bright sky^31, 32^, but they could also be identified as brightly reflecting objects against the darker foliage. How such bright targets are detected and tracked is currently not known.

### Behavioral relevance

TSDN responses to simple stimuli, such as the data shown in Figure 1-2, suggested that they should respond to reconstructed target pursuits. Indeed, the target sizes experienced during pursuit (Fig. 4C) matched the TSDN size tuning (Fig. 2A), as did the speeds (Fig. 2B, 4D). The target’s location in the visual field, however, was more equatorial (Fig. 4A) than the location of the receptive fields (Fig. 1, S1). This could be because the location of the target image (Fig. 4A) was reconstructed from the hoverfly heading (Fig. S2), which was based on the hoverfly’s location in the next time instance^33^, rather than on the hoverfly’s body orientation, and it did not include information about the head’s angle relative to the body.

Nevertheless, the predicted spike probability based on each neuron’s receptive field, directionality, and size tuning (Fig. 5, 6, S4) matched the observed spike probability well (Fig. 6H, 7, S4C), especially considering that we included no information about e.g. target trajectory history, nor its speed (Fig. 2C). Indeed, stimulus history is important for the appropriate encoding of naturalistic stimuli, as shown in e.g. macaque photoreceptors^48^. What was striking, however, was that the predicted spike probability, while correlated, was lower than the observed spike probability (Fig. S5). This suggests that the TSDNs respond substantially differently to simple stimuli, compared with those that are more naturalistic. Similarly, spider mechanoreceptors respond stronger to naturalistic stimuli^49^, as do *Drosophila* photoreceptors^50^.

We defined observed spike probability as the fraction of trials that generated at least one spike to any given frame within the reconstructed pursuits (Fig. 5). We found that the observed spike probability in individual neurons was relatively high (Fig. 9), with a median of 0.17, often reaching values of 1 (Fig. S5). In contrast, the population spike probability, which we defined as the fraction of neurons that had an observed spike probability of at least 0.5 to a given frame, was much lower (median 0.04, Fig. 9, S6D-E). This shows that not only was there a larger variation between TSDNs than within an individual TSDN, but only a few neurons responded to each part of each pursuit (Fig. 8A). Indeed, sparse population codes are common in sensory neurons, as shown in e.g. mouse primary visual cortex neurons in response to naturalistic stimuli^51^. Similarly, when ganglion cells are stimulated with dynamic stimuli, where the direction, velocity and luminance constantly change, the population response has higher information content than single cells do^52^. Furthermore, while simple stimuli (Fig. 1) suggested that we recorded from a homogenous population of neurons (Fig. S1), the responses to naturalistic stimuli highlighted that they were in fact rather heterogenous (Fig. 8, 9). This highlights the need for using more naturalistic stimuli when characterizing neurons.

### Neural coding in sensorimotor transformation

Descending neurons are incredibly important in their sensorimotor transformation role, as they constitute only a small subset (about 1%) of the central nervous systems of invertebrate and vertebrates alike, yet have to compress all the information from the brain into action commands projected to the body’s motor circuits^28, 53, 54^. An advantage of studying descending neurons in insects is that they are accessible, and sometimes it is even possible to record from them in behaving animals (e.g. Refs. ^55, 56^).

In insects, the descending neurons likely synapse with pre-motor or motor neurons that control behavioral output, such as e.g. movement of the wings, the neck or the legs^27, 28^. The descending neurons are likely to primarily act as initiators and modulators of the motor circuits that are located in the thoracic ganglia^57^. In this respect it is interesting to see how differently the individual TSDNs responded to reconstructed pursuits (compare e.g. Fig. 6G, S4B). Indeed, if we consider TSDNs in terms of behavioral output, rather than in terms of their responses to sensory input, the large variation could be beneficial. For example, one neuron may modulate head turning responses, which are often faster than the body’s response^58^, another TSDN might modulate abdominal ruddering^59^, and a third the wing beat amplitude^60^. In addition, for each of these body parts the descending neurons need to provide information about the direction and amplitude of the modulation. In the future, it will therefore be important to reconstruct the morphology of TSDNs, as done in dragonflies^22, 23^, as the location of their output synapses could inform us about their behavioral context. In addition, recording from TSDNs in behaving animals could substantially help us answer this question.

## Acknowledgements

We thank the Botanic Gardens of Adelaide for their ongoing support. This research was funded by the US Air Force Office of Scientific Research (AFOSR, FA9550-19-1-0294), the Australian Research Council (ARC, DP210100740, DP230100006 and FT180100289) and the EPSRC (EP/S030964/1 and EP/V052241/1).

