## Supplementary material for "Neural responses to reconstructed target pursuits": Supp Info

### Supplementary Figure Legends

**Figure S1. TSDNs belong to two clusters.** A) The optimal number of TSDN receptive field clusters after using *evalclusters* in Matlab. "All" includes each neuron's receptive field center location along the horizontal (X) and the vertical axis, its receptive field width, height, preferred direction (PD) and directionality index. We estimated the optimal cluster number under four more conditions, where "X + PD" used only the horizontal receptive field center location and the preferred direction, "All - X - PD" used all parameters except the horizontal receptive field center location and the preferred direction, "All - X" used using all parameters except the horizontal receptive field center location, and "All - PD" used all parameters except the preferred direction. B) The two clusters generated using the All condition. The data are color coded to show the receptive field centers of clusters 1 and 2, and their centroids.

**Figure S2. The conversion of 3D target pursuits into 2D representation for electrophysiology.** A) We defined the position of the bead relative to the hoverfly by subtracting the 3D hoverfly position from the bead position at each point in time. This allowed us to situate the hoverfly in the 3D origin (0, 0, 0). B) We determined the angle ($\alpha$) between the hoverfly flight heading ($H_{flight}$, black arrow) and a fixed heading ($H_{fixed}$, dashed magenta arrow) to the center of the screen, using the law of cosine. The original vector to the target is defined by $r$, and its rotated location by $r'$ (($x'_B$, $y'_B$, $z'_B$), magenta). C) For each frame we rotate the bead position by $\alpha$ using Rodrigues' formula, where $\vec{k}$ is the unit vector in the direction of the rotation axis, $r$ is the original vector to the target, and $r'$ its rotated location. D) We computed the screen position ($r'_{screen}$) of the bead in each frame, where $z_{screen}$ is the distance between the immobilized hoverfly and the center of the screen (6.5 cm). E) The size of the bead in each frame was calculated using the length of the vectors $r'_{screen}$ and $r'$, and the physical bead size ($w$), and converted to pixels.

**Figure S3. TSDN inter-spike interval distribution follows a gamma function better than a Poisson function.** The root mean squared error (RMSE) between a gamma or a Poisson function and the distribution of TSDN interspike intervals (ISI), as quantified in each TSDN (N = 27). The median RMSE values were 0.09 for the gamma function and 0.34 for Poisson function. The black dots show the example neuron in Figure 1A, B, 5 and 6. Significance was investigated using the Wilcoxon test: ****$P < 0.0001$

**Figure S4. The observed and predicted spike probability in a highly responsive TSDN.** A) The predicted spike probability for an example TSDN to the six different target trajectories. B) The observed spike probability for the same TSDN to the same six target trajectories. C) The correlation between predicted and observed spike probability to the six target trajectories.

**Figure S5. The maximum predicted spike probability is significantly lower than the maximum observed spike probability.** The maximum observed and predicted spike probability measured from each TSDN (N = 27), from any of the six pursuits. The median values were 1 for the observed and 0.46 for the predicted spike probability. Significance was investigated using the Wilcoxon test: ****$P < 0.0001$.

**Figure S6. Calculation of population spike probability.** A) Each row in the raster plot shows the response of one TSDN to an example reconstructed pursuit (same as in Fig. 3, 5, 6). Each vertical line indicates that the TSDN in that row had an observed spike probability above 0.5 (see Fig. 5) for that frame. Yellow and orange arrows highlight the example neurons from Figure 6 and Figure S4. B) The green bars show the frames where any of the 27 TSDNs had an observed spike probability above 0.5. C) The same trajectory as reconstructed on the screen, where green, filled symbols correspond to the green coloring in panel B. D) A magnification of the dashed box shown in panel A. Pale blue highlights frame 270 (5 neurons out of 27 responding) and dark blue frame 281 (3 neurons out of 27 responding). E) Population spike probability, which is the fraction of TSDNs that responded to a given frame (0.19 for frame 270, 0.11 for frame 281). F) The same data as in panel E, but shown as a reconstructed trajectory on the screen, and color coded according to population spike probability. The colored arrows highlight frames 270 and 281.

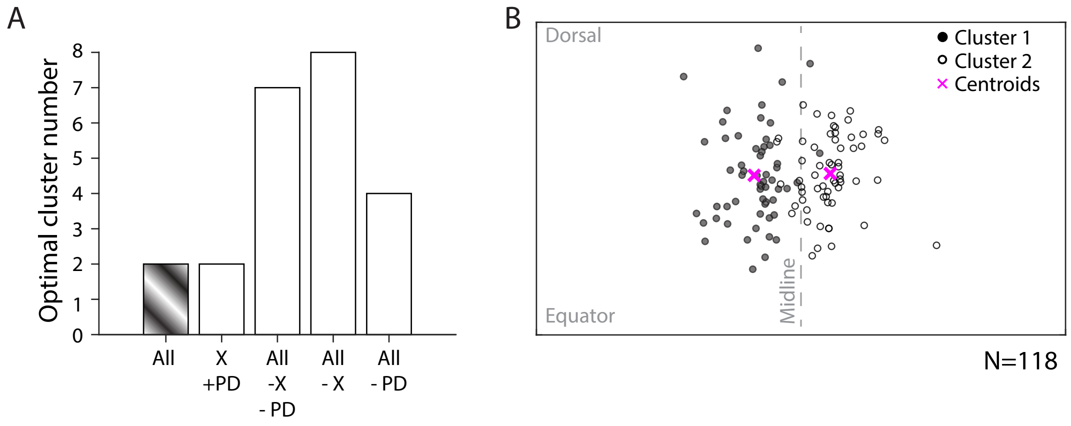
A
Optimal cluster number
8
7
6
5
4
3
2
1
0
All
X
+PD
All
-X
- PD
All
- X
All
- PD
B
Dorsal
Equator
Midline
Cluster 1
Cluster 2
Centroids
N=118

Figure S1

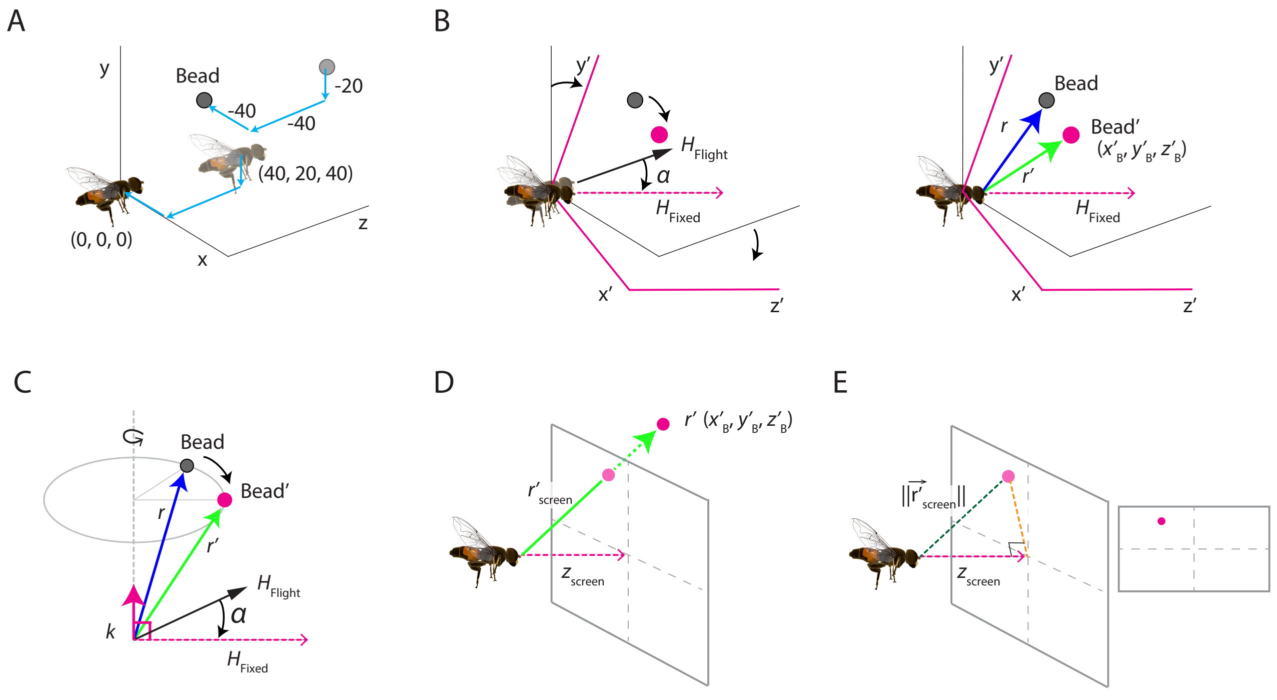
A
y
Bead
-20
-40
-40
(40, 20, 40)
(0, 0, 0)
x
z
B
y′
H Flight
α
H Fixed
x′
z′
y′
Bead
r
Bead′
(x′B, y′B, z′B)
r′
H Fixed
x′
z′
C
Bead
Bead′
r
r′
H Flight
α
k
H Fixed
D
r′ (x′B, y′B, z′B)
r′ screen
z screen
E
||r′ screen||
z screen

Figure S2

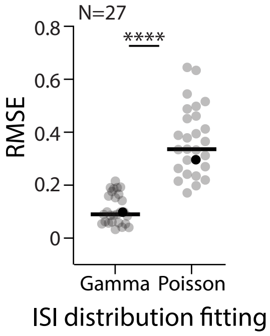
N=27
****
0.8
0.6
0.4
0.2
0
RMSE
Gamma
Poisson
ISI distribution fitting

Figure S3

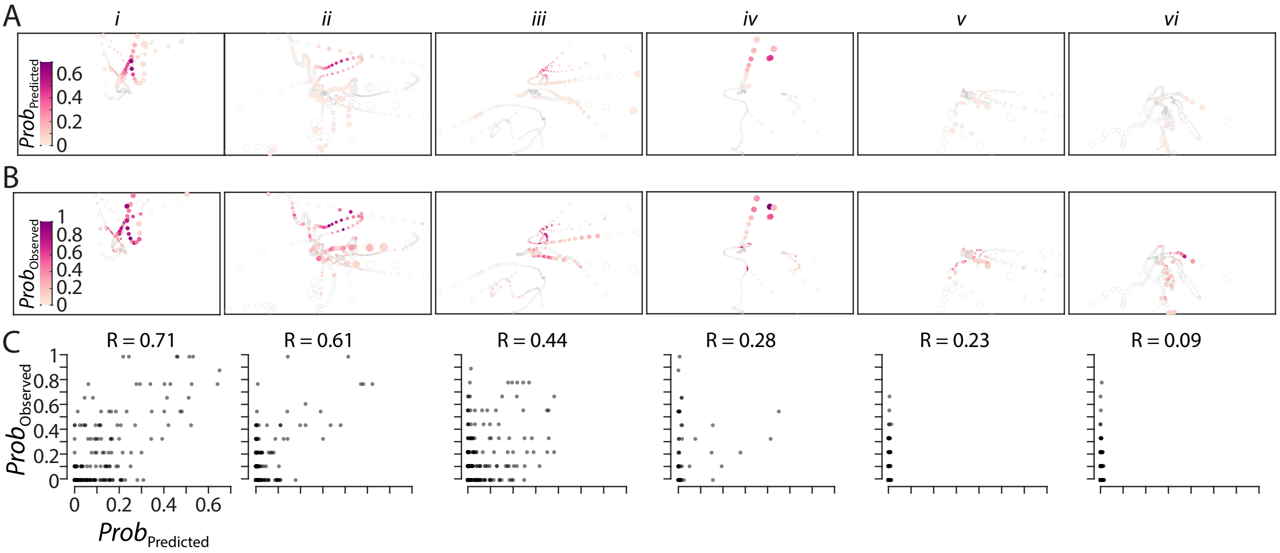

A
B
C
i
ii
iii
iv
v
vi
Prob_Predicted
0.6
0.4
0.2
0
Prob_Observed
1
0.8
0.6
0.4
0.2
0
R = 0.71
R = 0.61
R = 0.44
R = 0.28
R = 0.23
R = 0.09


Figure S4

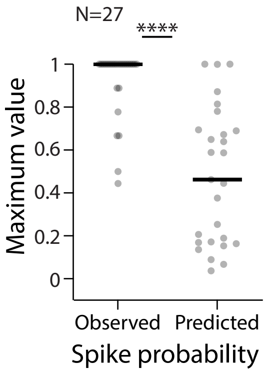

N=27
****
Maximum value
1
0.8
0.6
0.4
0.2
0
Observed
Predicted
Spike probability


Figure S5

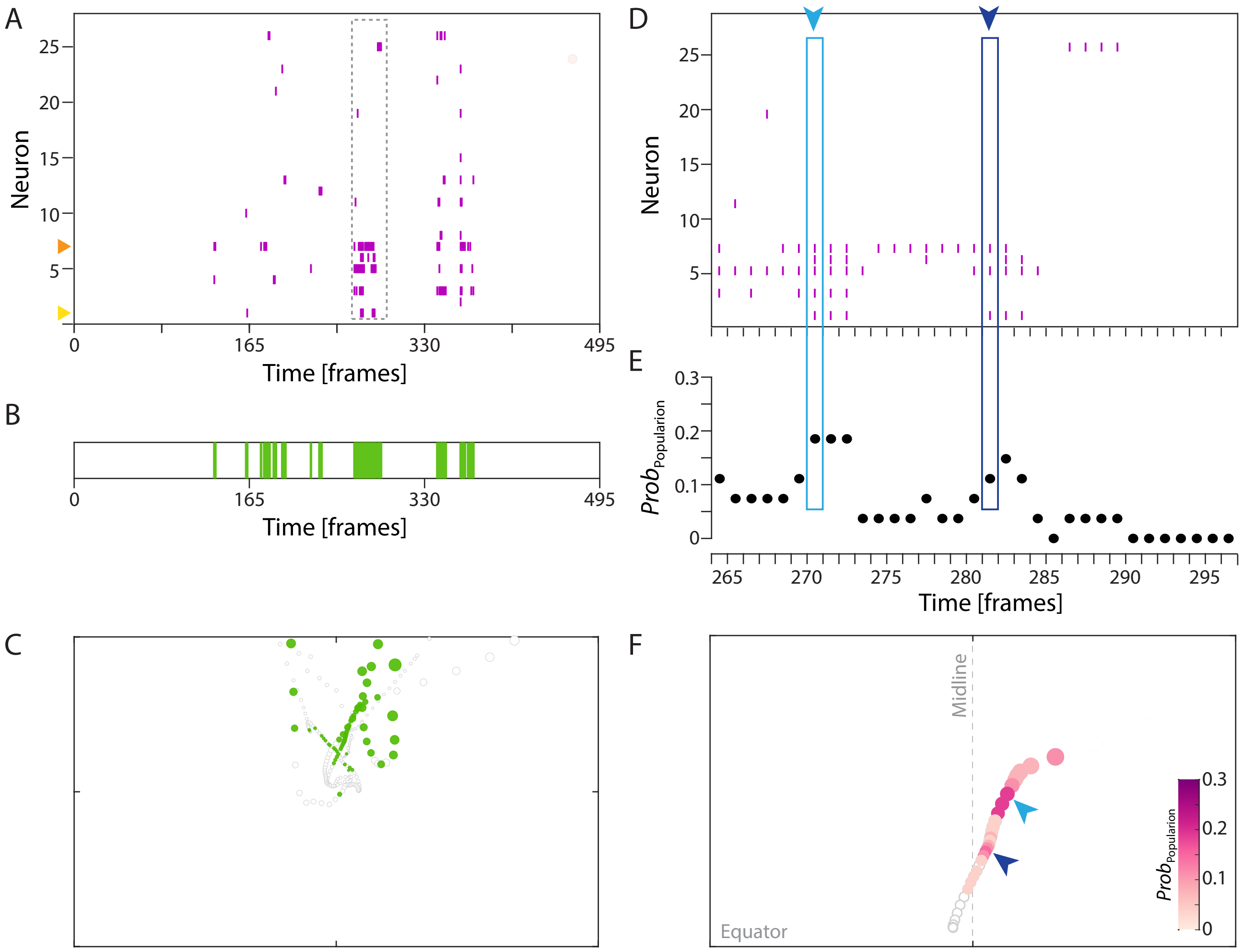
A
Neuron
25
20
15
10
5
0
165
330
495
Time [frames]
B
0
165
330
495
Time [frames]
C
D
Neuron
25
20
15
10
5
E
$Prob_{Popularion}$
0.3
0.2
0.1
0
265
270
275
280
285
290
295
Time [frames]
F
Midline
Equator
$Prob_{Popularion}$
0.3
0.2
0.1
0

Figure S6
